# Stochastic dispersal increases the rate of upstream spread: a case study with green crabs on the northwest Atlantic coast

**DOI:** 10.1101/188409

**Authors:** A. Gharouni, M.A. Barbeau, J. Chassé, L. Wang, J. Watmough

**Affiliations:** Department of Mathematics and Statistics, University of New Brunswick, Fredericton; Department of Biology, University of New Brunswick, Fredericton; Fisheries and Oceans Canada, Gulf Fisheries Centre, Moncton, NB

**Keywords:** Aquatic Invasive Species, Dispersal heterogeneity, Integro-difference equation, Larval dispersal, Spatial population model, Spread rate

## Abstract

Dispersal heterogeneity is an important process that can compensate for downstream advection, enabling aquatic organisms to persist or spread upstream. Our main focus was the effect of year-to-year variation in larval dispersal on invasion spread rate. We used the green crab, *Carcinus maenas*, as a case study. This species was first introduced over 200 years ago to the east coast of North America, and once established has maintained a relatively consistent spread rate against the dominant current. We used a stage-structured, integro-difference equation model that couples a demographic matrix for population growth and dispersal kernels for spread of individuals within a season. The kernel describing larval dispersal, the main dispersive stage, was mechanistically modeled to include both drift and settlement rate components. It was parameterized using a 3-dimensional hydrodynamic model of the Gulf of St Lawrence, which enabled us to incorporate larval behavior, namely vertical swimming. Dispersal heterogeneity was modeled at two temporal scales: within the larval period (months) and over the adult lifespan (years). The kernel models variation within the larval period. To model the variation among years, we allowed the kernel parameters to vary by year. Results indicated that when dispersal parameters vary with time, knowledge of the time-averaged dispersal process is insufficient for determining the upstream spread rate of the population. Rather upstream spread is possible over a number of years when incorporating the yearly variation, even when there are only a few “good years” featured by some upstream dispersal among many “bad years” featured by only downstream dispersal. Accounting for annual variations in dispersal in population models is important to enhance understanding of spatial dynamics and population spread rates. Our developed model also provides a good platform to link the modeling of larval behavior and demography to large-scale hydrodynamic models.

## Introduction

Understanding the mechanisms used by invasive species to spread against a dominant current in an aquatic environment remains an interesting problem today despite more than two decades of research. Upstream spread is related to the “drift paradox”, whereby populations persist even when subjected to continuous advection [1]. One hypothesis is that variability in flow direction (e.g., due to turbulence or tides) coupled with high reproductive rates is sufficient to compensate for the downstream loss of individuals [2–4]. In marine systems, we see flow variability due to widely different phenomena [5]. These can be grouped broadly into two relevant time scales: (i) the dispersal period (typically the larval stage) and (ii) the adult lifespan. Even when variability on the shorter (within year) scale is not sufficient to resolve the drift paradox, longer-scale variability (year-to-year) may still allow persistence and upstream spread [3]. Byers and Pringle [3] showed that for several simple models, dispersal strategies in which an adult releases larvae either over several years or several times each year increase the likelihood of population retention and spread against a dominant current. In the present study, we more closely examine this idea using a stage-structured integro-difference equation (IDE) model whose dispersal kernel more explicitly models larval dispersal in currents and allows for year-to-year fluctuations in both speed and variability of currents. We conclude that increased year-to-year variability in currents leads to increased spread rates against the dominant current. Moreover, basing dispersal parameters on mean flows underestimates spread rates.

The green crab (*Carcinus maenas*) is a particularly interesting case study because it is a high impact invasive species and is an effective disperser [6–8]. On the coast of New England and Maritime Canada, the northward spread rate appears fairly consistent over a large temporal scale (e.g., 180 km per decade over 120 years) and spatial scale (along the Gulf of Maine, the Atlantic coast of Nova Scotia and the southern Gulf of St Lawrence) [8,9], despite spreading through water bodies with different characteristics and the occurrence of multiple introductions [10]. Further, this invasion occurred against the dominant (southwest) flow [11,12]. Note that on a short time scale (when yearly measurements are available), the spread rate appears more punctuated, with little spread in some years and large spread in other years (e.g., [13]).

There is a need for more careful study of dispersal. Not only is it challenging to obtain good estimates of dispersal distances, past modeling studies have shown that spread rates are highly sensitive to the extent and frequency of the furthest dispersal distances [14] and that larval dispersal patterns can determine the population structure of an invasive species throughout its spatial domain [15]. In our context, we are focusing on dispersers moving through natural means against a dominant current. Note that accounting for anthropogenic movement could affect any model predictions. The multiple introductions of *C. maenas* in Newfoundland are an example of such movement [16]. In coastal marine species, the life stage with the greatest natural dispersal distances is typically the larval stage (or propagule stage for non-animal species) [17]. Therefore, it is important to determine the processes that influence larval dispersal, and incorporate them in a modeling framework (our work provides such a framework). These processes can be categorized into three groups [17]: (i) biological processes including production, growth, development, and survival of larvae; (ii) physical processes such as currents and turbulence; and (iii) behaviors, such as larval vertical swimming, that link the two.

In summary, the main goal of our paper was to examine the effect of year-to-year variability in dispersal on the spread rate of invasive species against a dominant flow, building on previous modeling studies [3,9,18,19]. Byers and Pringle [3] examined the importance of variability in water currents on retention and, based on the results of Pachepsky et al. [19], inferred that retention and spread were linked. We focused here on the effect of variability in the dispersal kernel (which incorporates advection, diffusion and larval behavior) on spread rate (and not only the likelihood of spread). We used a stage-structured IDE model extended from the model developed by Gharouni et al. [9] with the addition of a mechanistic and stochastic dispersal component to investigate the northward spread rate (i.e., against the dominant current) of the green crab on the east coast of North America. Gharouni et al. [9] investigated the compensatory relationship between demographic and dispersal parameters for a given spread rate using a linear and density-independent, deterministic, and stage-structured IDE model. Note that a compensatory relationship between dispersal and reproduction has been established since Fisher’s seminal work [18]. Here, we confirmed that a similar relationship persists in the IDE framework even when incorporating density dependence and more complex dispersal characterizations than previously done (see Discussion). Although we used the green crab invasion as a case study, our model can be modified for any aquatic species of interest. Our model has two components, a demographic projection matrix describing reproduction and transitions between life stages (first-year juvenile, second-year juvenile and adult), and probability distributions (kernels) modeling dispersal of each stage. For the larval dispersal kernel, we used a mechanistic and flexible approach following Pachepsky et al. [4]. We first used a deterministic version of the model to verify the relationship between the spread rate and the dispersal and demographic parameters (net rate of displacement, diffusion and recruitment). Note that we used the term “net rate of displacement” rather than “advection”, because behavior and various other processes in addition to passive advection influence how far a disperser is displaced. We then allowed dispersal (specifically, the net rate of displacement and diffusion coefficient of larvae) to vary yearly by parameterizing the dispersal kernels using a 3-dimensional hydrodynamic model of the Gulf of St Lawrence [20,21]. The same hydrodynamic model was used for several other biological dispersal studies [22–25]. Our IDE framework is well-suited to study the effect of heterogeneous dispersal on invasion spread rates. The within-year variability in dispersal distances is reflected in the shape of the larval dispersal kernel, which incorporates components for drift and settlement rate and can be affected by larval behavior. Year-to-year variability is modeled by allowing the parameters of the kernel to vary stochastically.

## Methods

### Model framework

Our model consists of a system of IDEs describing the population dynamics of three stages of female crabs: first-year female juveniles, second-year female juveniles, and mature female adults. This stage structure captures the important difference in time scales between the maturation of the juveniles and dispersal of larvae [9]. The larval stage, whose development and dispersal occurs offshore on a shorter time scale, is implicitly included in fecundity and dispersal components of the model [8]. The general form of the model is

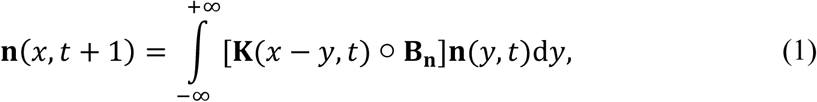

where **n** = (*n*_1_, *n*_2_, *n*_3_) denotes the vector of abundances of each stage at discrete time (*t*) and continuous position (*x*). **B**_**n**_ and **K** denote, respectively, the population projection matrix and the dispersal matrix, and are defined as

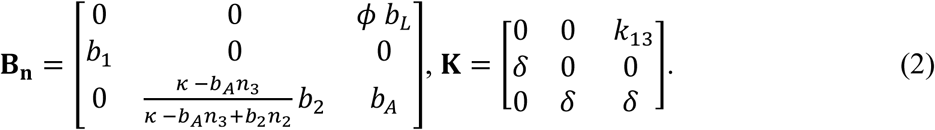

The projection matrix (**B**_**n**_) contains the demographic components of the model. The parameters include the average number of female eggs produced per mature female per year (*ϕ*), the probability that a given egg survives to become a settled first-year juvenile (*b*_*L*_), the survival probabilities for first and second year juveniles (*b*_1_ and *b*_2_, respectively) and the survival probability of adults (*b*_*A*_). The fraction of second-year female juveniles that survive to become adults (row 3, column 2 entry of **B**_**n**_) is assumed to be density dependent. This density dependence is a refinement of the model developed by Gharouni et al. [9], which was based solely on the linearized model; the refinement was done to make the density dependence explicit, in the form of competition for space between year-two juveniles and adults. Specifically, we assumed that there are *κ* sites available for adults to occupy, that surviving female adults from the previous year, *b*_*A*_*n*_*3*_, continue to occupy their sites, and that second-year female juveniles then compete for the remaining spaces. Thus, a fraction, 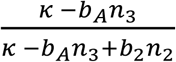, of the *b*_2_*n*_2_ maturing year-two juveniles obtain sites. Note that if *n*_2_ is small relative to the number of remaining sites, then the number surviving approaches *b*_2_*n*_2_ and if *n*_2_ is large, the number of survivors approaches the number of remaining sites.

Although our main interest is the effect of annual variations in dispersal distances, we first considered the case where the dispersal matrix, **K**, has no year-to-year variations. The larval dispersal kernel, *k*_13_, models the dispersal of the current year’s offspring that survive to become first-year juvenile recruits. The time scale of the larval dispersal process directed our characterization of time as being discrete and in having a model time-step of one year. We assumed that spread is driven by larval dispersal, so dispersal in other stages is modeled by the Dirac delta, *δ*, which represents no movement. We used a mechanistic larval dispersal kernel for *k*_13_ following Pachepsky et al. [4]:

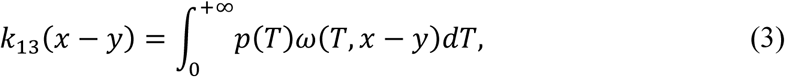

where *p(T)* is the probability a given larva settles at time *T*, conditional on it having survived the dispersal process, and *ω*(*T,x* – *y*) is the probability distribution of positions of dispersing larvae at a time *T* following release. This approach enabled us to describe the dispersal process in more detail, incorporating known settlement windows and drift components, than using a simple Normal or Laplace distribution as in the model developed by Gharouni et al. [9]. For our paper, we assumed a uniform distribution of settling rates

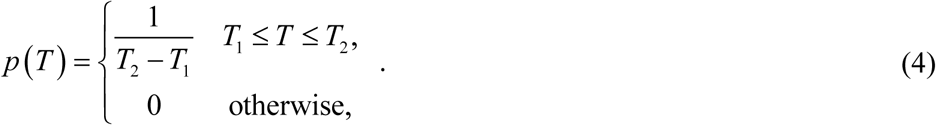

That is, the duration of the larval pelagic period ranges from *T*_*1*_ (earliest settlement) to *T*_*2*_ (latest settlement). We modeled larval drift as a normal distribution:

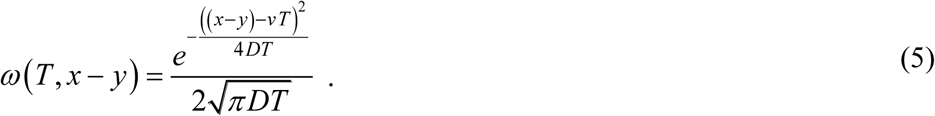

Such larval drift is a common assumption for marine species with a pelagic larval stage (e.g., [26,27]). More specifically, we assumed larvae drift with mean displacement *μ* = *vT* and standard deviation 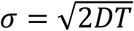, where *v* is the net rate of displacement and *D* is the diffusion coefficient. Note that the settlement rate and drift components could take a number of different shapes depending on the study organism, as discussed later. We refer to Eqs. (1-5) as our deterministic model. Below (in the section “Stochastic model spread rates”), we introduced a time dependence into the dispersal matrix **K** by allowing *v* and *D* to vary with year; we refer to Eqs. (1-5) with year-to-year varying *v* and *D* as our stochastic model.

In Fig. 1, we illustrated the shape of kernel *k*_*13*_ and its dependence on *v* and *D*. The different curves were numerically computed using Eqs. (3-5) with different pairs of *v* (net rate of displacement) and *D* (diffusion coefficient). Note that for our choice of *p* and *ω* (the settlement and drift components), *k*_*13*_ does not have an explicit representation in terms of basic functions. Also, note that we orient our coastline so that positive displacements are against the dominant current, and *v* is generally negative. Since the dominant current is southward in our selected study region, positive spread rates represent a northward invasion or range expansion. In Fig. 1, a small fraction of individuals move northward even with a southward net rate of displacement (negative *v*) [4]. The shape of the kernel changes with different values of *v* and *D*, because of the interplay between the modifying effect of the settling rate function and smoothing effect of the integration. Unlike the shifted Normal distribution, which is a rigid shift (no change in shape), this kernel shifts, flattens and appears to approach a uniform distribution as *v* increases.

**Fig. 1.**
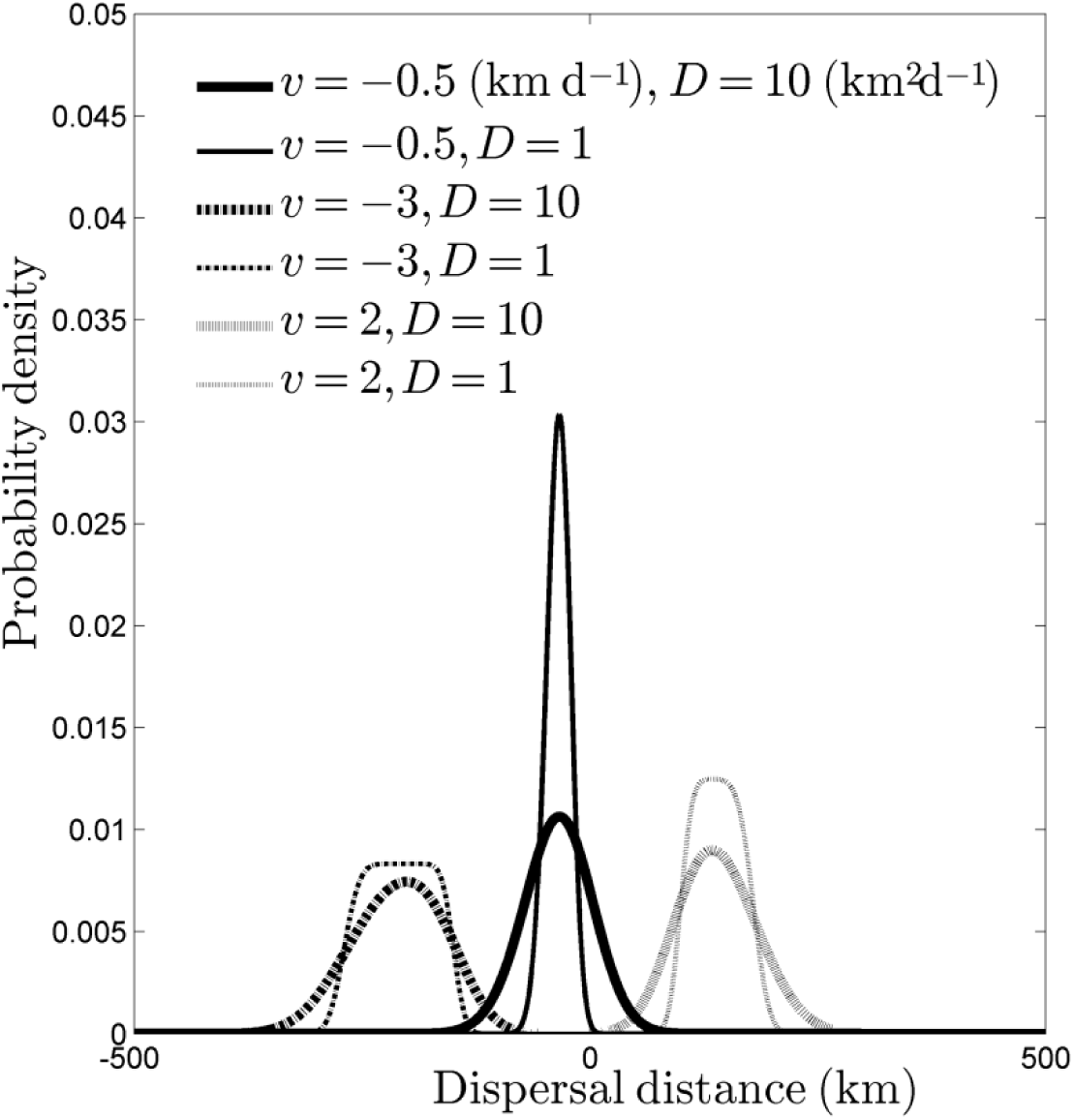
Representations of the larval dispersal kernel *k*_*13*_ (Eqs. 3-5) with different values for the net rate of displacement *v* (km d^−1^) and diffusion coefficient *D* (km^2^ d^−1^).

For this graph, *T*_1_ = 50 d and *T*_2_ = 90 d. Note how the kernel shifts, flattens, and appears to approach uniform distribution as the magnitude of *v* increases.

For deterministic models in the general form of Eq. (1), it is conjectured that solutions converge to a travelling wave solution with speed *c*\* provided certain conditions are met [28], where *c*\* is the speed of the stable travelling wave solution of the linearization of the model near the trivial equilibrium (**n** = **0**). The conditions of the conjecture are that (i) the initial conditions are bounded, positive and zero outside a closed interval, (ii) the leading eigenvalue of **B**_0_ is larger than one, (iii) **0** ≤ **B**_**n**_ **n** ≤ **B**_**0**_ **n**, and (iv) the entries of **K** have moment generating functions. We have verified that these conditions hold for our deterministic model. Linearization of our model, Eqs. (1-5), in the vicinity of **n** = **0** leads to using **B**_**0**_ in Eq. (1) in the analysis. Note that **B**_**0**_ here is identical to the demographic matrix, **B**, used in the linear (density independent) model developed by Gharouni et al. [9]. The computations of *c*\* follow the procedures detailed in Gharouni et al. [9], with the addition that the moment generating function for *k*_*13*_ is computed numerically. Similar to the results found in Gharouni et al. [9], there is a threshold in the *v*-*D* parameter plane such that if *v* and *D* lie below this threshold, there is no northward travelling wave solution (*c*\* < 0; see Fig. 2). If *v* and *D* are above this threshold, *c*\* ≥ 0 and there is a northward travelling wave solution. Thus, following the terminology introduced by Caswell et al. [29], we refer to a pair of *v* and *D* as a “good-year” pair if they lie above the threshold and a “bad-year” pair otherwise.

**Fig. 2.**
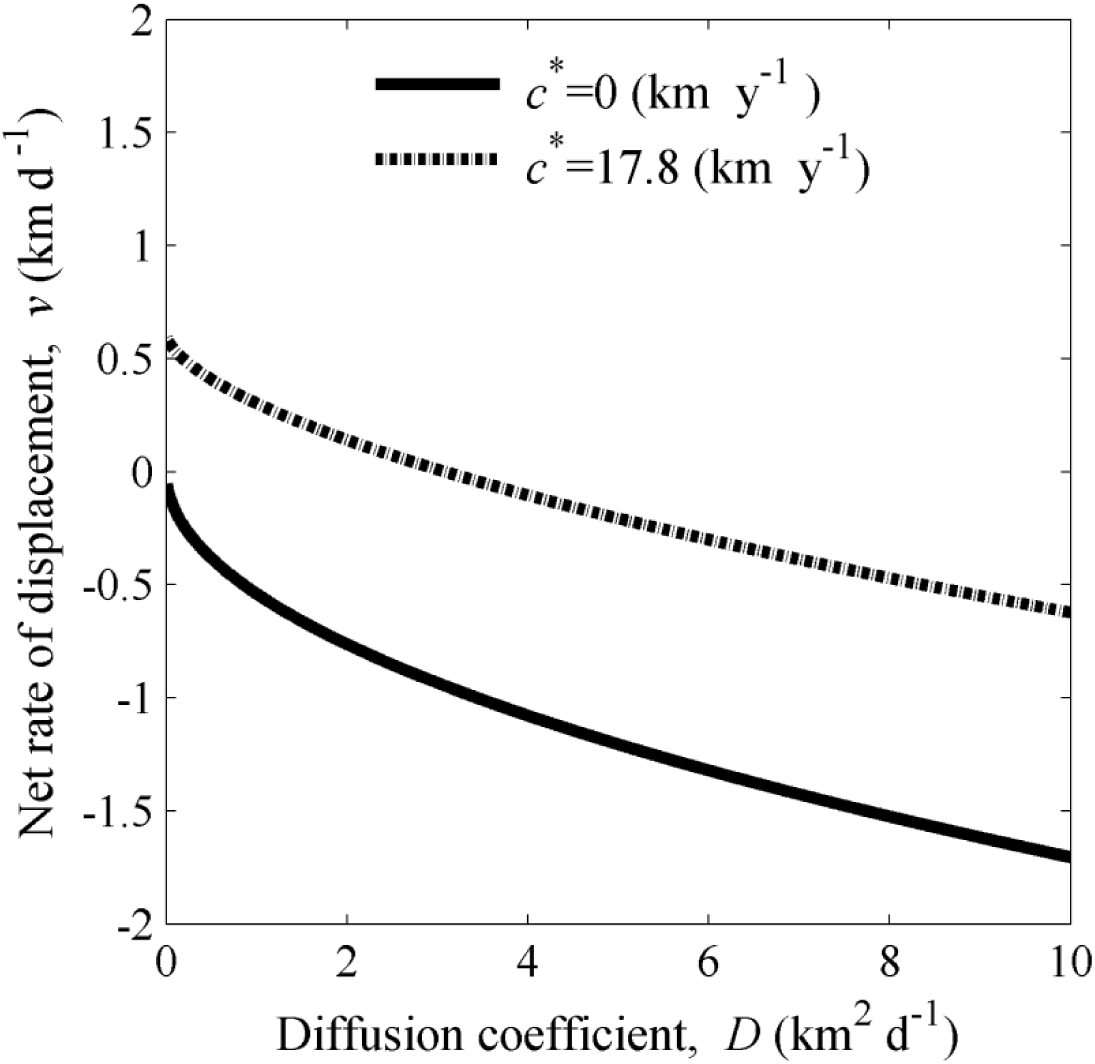
Contours for spread rate *c*\* against a dominant current.

The solid line shows the threshold for northward spread, i.e., the relationship between net rate of displacement *v* and diffusion coefficient *D* suggested by our model (Eqs. 1-5) beyond which northward spread can occur (*c*\* ≥ 0 km y^−1^). A pair of values of *v* and *D* is referred as a “good-year” pair if it lies above this threshold, otherwise the pair is referred as a “bad-year” pair. The dash-dot line is the contour for *c*\*=17.8 km y^−1^, which is the estimated northward spread rate of the northern lineage of the green crab in the Northumberland Strait, Canada [9]. The other model parameters were fixed as adult survival probability *b*_*A*_ = 0.8, recruitment rate *r* = 23 adult female offspring per female adult per year, and the start and end of the larval settlement period *T*_1_ = 50 and *T*_2_ = 90 days.

### Parameter estimation

Estimates for all parameters (*r*, *b*_*A*_, *T*_*1*_, *T*_*2*_, *v*, *D*) were made using independent sources. Additionally, we also estimated the northward spread rate (see below) from another independent data set. Details of these estimates are discussed below, and sources and further details can be found in Gharouni et al. [9]. The annual survival rate of female adults was estimated as *b*_*A*_=*0.8* based on a total lifespan of 6 years [30]. Following [9], we defined the adult recruitment rate as *r*=*ϕb*_*L*_*b*_*1*_*b*_*2*_, and estimated it as *r*=*23* adult female offspring per female adult per year. Without loss of generality, we assumed *κ*=1 in our simulations; this is equivalent to measuring populations as fractions of the available sites per km. Green crab larvae spend between 50 and 90 days in coastal waters and then return inshore to settle [8]; thus, we used *T*_*1*_=50 d, *T*_*2*_=90 d for all calculations and simulations.

We used an independently estimated northward spreading speed of *c*\*=17.8 km y^−1^ for the northern lineage of green crabs throughout Northumberland Strait up to the Kouchibouguac Lagoon (New Brunswick) from 1994 to 2013 (data on green crab sightings from Fisheries and Oceans Canada; regression analysis of northern-most sighting in a given year, representing the invasion front, detailed in [9]). We chose the Northumberland Strait as the region for our case study because of the high quality of the field data for the green crab invasion.

To estimate values for the dispersal parameters (*v* and *D* in *k*_13_) for a given year, we used a hydrodynamic model consisting of (i) a particle-tracking, individual-based model run within (ii) an ocean circulation model based on the Nucleus for European Modeling of the Ocean (NEMO) system described in detail in Brickman and Drozdowski [20] and Lavoie et al. [21]. The modeling system is based on the ocean code OPA version 9.0 [31]. The domain of the ocean circulation model includes the Gulf of St. Lawrence, Scotian Shelf and Gulf of Maine. The horizontal resolution is 1/12° in latitude and longitude, and the vertical resolution has 46 layers of variable thickness (from near the water surface to about 250 m in depth). This ocean circulation model is prognostic, allowing for advection-diffusion of the temperature and salinity fields, which are only constrained through open boundary conditions, freshwater runoff and surface forcing. It also includes tidal forcing. This ocean circulation model drove a bio-physical model (the individual-based model) that can simulate advective dispersal of “particles” representing green crab larvae in nearshore flow and hydrodynamic fields. The particle-tracking model enabled us to virtually release and follow larvae that have behavior (such as vertical swimming) under variable environmental conditions. Because of data storage constraints, the physical velocity fields from the ocean model were outputted as daily averages, but they were interpolated to hourly values to drive the particle-tracking model offline. The small-scale diffusion coefficient in the particle tracking model was set at 25 m^2^ s^−1^ following the work of Chassé and Miller [32] and Hrycik et al. [33] to represent within-day variation such as tidal stirring, wind waves, tides, etc. The depths within which the larvae were virtually swimming was set between 20 m and 30 m in the Northumberland Strait; specifically, this represented larvae swimming up to 20 m at nighttime and down to 30 m in daytime [34,35].

To obtain sample larval dispersal kernels, we simulated the release of cohorts of larvae from 8 selected locations along the mainland coast of the Northumberland Strait using the hydrodynamic model for each year from 2007 through to 2012 (Fig. S.1). The release dates of the larvae within a year were daily from July 1^st^ to 31^st^, and every hour during each day of the release period (to approximate the spawning period of green crabs in this region) [8]. The procedure of virtual releasing and tracking was repeated 5 times for a given location and year to better capture environmental variability. The trajectories of the particles were tracked for 90 days from the date they were released. Since, based on our settling rate function *p(T)* (Eq. 4), particles can settle as early as 50 days after release, we recorded their location every day between day 50 and 90. The output locations (latitude and longitude) were projected onto a defined 1-dimensional coastline. The coastline was modeled by a straight line (dashed line in Fig. S.1). The dispersal displacement for a given particle is defined as the distance between its projected locations at the release time and settling time. This represents the dispersal displacement of a larva from the source.

For each of the 8 release locations and each of the 6 years, we obtained a frequency distribution of dispersal displacements of larvae (i.e., 48 simulated dispersal kernels; Fig. S.2.). We then used a maximum likelihood statistical procedure (*fminsearch* function in Matlab^®^ [36]; Math Works) to fit our theoretical dispersal kernel (Eqs. 3-5) to each simulated dispersal kernel to estimate the dispersal parameters, *v* and *D*. Thus, we obtained a pair of *v* and *D* estimates for each release location-year combination, for a total of 48 estimated pairs of (*v,D*) (Fig. S.2 and Table S.1 in the online supplement). These estimated pairs of dispersal parameters were used to study the effect of year-to-year larval dispersal variability on the spread rate.

### Stochastic model spread rates

The linear conjecture and accompanying spread rate formulae cannot be applied to our stochastic model. Hence, we used numerical simulations to determine the mean spread rates with year-to-year variations in dispersal. Simulations were carried out in Matlab^®^ [36]: for each simulation, the model was run over 30 time-steps with initial conditions symmetric about the point *x* = 0 on the domain −3000 km ≤ *x* ≤ 3000 km. The integral in Eq. (1) was approximated by using the trapezoidal method (via the Matlab function *trapz*). The spatial domain was chosen sufficiently large so that a travelling wave developed before reaching the boundary. The location of the forward wave front (rightward in the 1-D spatial domain) was defined to be the right-most point for which the adult population density was above *κ*/2. The spread rate for a simulated solution to the stochastic model was then taken to be the slope of a linear regression of the wave-front locations.

To evaluate the effect of stochastic dispersal on the spread rate, *c*\*, we did the following analysis.

i. We chose 100 sample time-sequences of size 30 by randomly sampling with replacement from the 48 estimated pairs of (*v,D*) (Table S.1).
ii. For each sample, we ran the model using the sequence of pairs of (*v,D*). This resulted in a sample-specific “stochastic spread rate” which is denoted by *c*_s_.
iii. We computed an “averaged (*v,D*)” for each sample by calculating the arithmetic mean of the 30 kernels corresponding to each time-sequence of pairs of (*v,D*) obtained in step (i), and fitting the theoretical dispersal kernel (Eqs. 3-5) to the result using the *nlinfit()* function in Matlab^®^. The spread rate resulting from this averaged (*v,D*) is referred to as the “averaged spread rate” and denoted by *c*_a_.
iv. We computed a “grand-averaged” estimate of (*v,D*) from the 48 estimated pairs of (*v,D*) (Table S.1) by the same method of step (iii). The spread rate resulting from the grand-averaged (*v,D*) is referred to as the “grand-averaged spread rate” and denoted by *c*_g_.

The same initial conditions were used for all model runs. We used a developed travelling wave, because our focus is on the asymptotic spread rate and not the transient dynamics. To obtain the developed travelling wave, we ran the model for 20 time steps using the grand-averaged estimate of (*v,D*) with an initial population density of zero for first-year and second-year juveniles, *n*_*1*_(*x,0*)= *n*_*2*_(*x,0*)=*0*, and of one for the adults, *n*_*3*_(*x,0*)=1 for *x* in [-*2,2*] and zero otherwise. 20 time-steps was observed to be sufficiently long for a travelling wave to develop.

After doing (i) above, we found that all 48 pairs of (*v,D*) obtained from the hydrodynamic model for the Northumberland Strait between 2007 and 2012 were “good-year” pairs. Since we were interested in gaining insight for scenarios where northward spread did not occur every year, we obtained a second set of 100 samples by simply subtracting 3 km d^−1^ from *v* in each of the 100 samples obtained in step (i) above. This resulted in a mixture of good and bad years in each time-sequence, to reflect scenarios of other coastlines (such as the Atlantic coast of Nova Scotia, based on our preliminary examination of other coastlines) (see also [12]). Then, we repeated steps (ii), (iii), and (iv) with these shifted (*v,D*) pairs. Spread rates resulting from the shifted pairs are denoted by 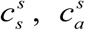, and 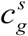 for shifted stochastic, shifted averaged, and shifted grand-averaged spread rates, respectively.

Finally, we compared the stochastic, averaged and grand-averaged spread rates by computing the differences *c*_*s*_-*c*_*g*_, *c*_*s*_-*c*_*a*_, and *c*_*a*_-*c*_*g*_, as well as 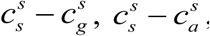, and 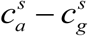.

## Results

Using *c*\*=17.8 km y^−1^ for an estimated mean spread rate against the dominant current for green crabs [9], our deterministic model (Eqs. 1-5) provided a set of feasible net rates of displacement, *v*, and diffusion coefficients, *D*, for a given recruitment rate, *r* = *ϕb*_*L*_*b*_1_*b*_2_(Fig. 2 and 3). Moreover, setting *c*\*=0 km y^−1^ (termed the critical spread rate) provided a threshold for dispersal parameters above which the population spreads northward and below which the population retreats southward (Fig. 2). When setting the recruitment rate to a reasonable estimate from field data for the green crab (*r* = 23 adult female offspring per female per year [9]) for these two situations (the observed and the critical spread rates), the curve representing the feasible set of *v* and *D* is concave and decreasing in the *v-D* plane (Fig. 2). This indicates a compensatory relationship between the diffusion coefficient and the net rate of displacement. Specifically, when a cohort of larvae experiences a southward net rate of displacement (*v*<0) increased diffusion can compensate, leading to spread against the current. In contrast, in years when currents are reversed, resulting in a northward net rate of displacement (*v*>*0*), the diffusion coefficient can be very low. When the recruitment rate, *r*, is increased, the feasible values of *v* and *D* can be reduced for a given spread rate (Fig. 3). Thus, in keeping with the theory developed by Fisher [18] and Byers and Pringle [3], high southward currents can be compensated for by either increased *D*, increased *r*, or a combination of the two.

**Fig. 3.**
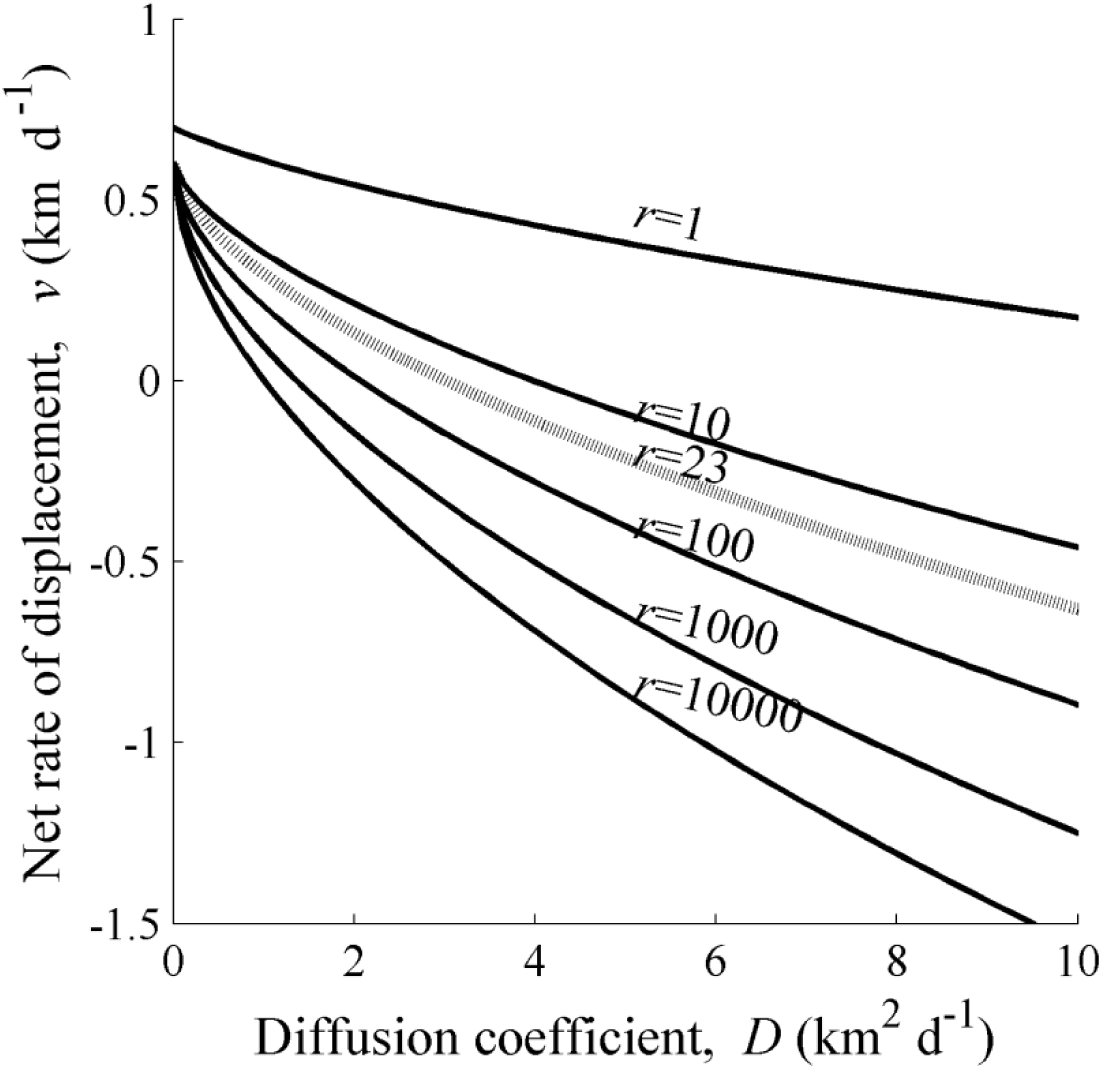
Level curves at spread rate *c*\*=17.8 km y^−1^ for different values of recruitment rate *r*.

The dotted curve is the *c*\*=17.8 contour for *r* = 23 (adult female offspring per female per year) and is the estimate for recruitment rate for green crabs *Carcinus maenas* on the northwest Atlantic coast (based on biological information compiled in Gharouni et al. [9]). If the values of *v* and *D* lie below one of these curves, then the deterministic model predicts slower or no northward spread (against the dominant current) for that particular value of *r*. The other model parameters were fixed as *b*_*A*_ = 0.8, and *T*_1_ = 50 and *T*_2_ = 90 days. Note that the dotted curve in this figure is the same as the dotted curve in Fig. 2.

Environmental stochasticity can also lead to spread against the dominant current. The set of simulations seeded with both good years and bad years resulted in northward spread even when the averaged dispersal kernels did not support northward spread (Fig. 4c). In these simulations, almost all spread rates using averaged kernels were smaller than spread rates using stochastic kernels (99 out of the 100 samples; middle boxplot in Fig. 4d). In addition, spread rates using averaged kernels were negative for all 100 samples, while spread rates using stochastic kernels were positive for 89 out of the 100 samples (Fig. 4c). Similarly, in simulations run with only good years, the spread rates from the model using stochastic kernels were all higher than those from the model using the averaged kernels (Fig. 4a and b). As expected, the comparison of the model using the sample-averaged kernel to that using the grand-averaged kernel (right boxplots in Fig. 4b and d) did not differ much, which is an indication that sampling (with replacement) 30 out of the 48 kernels is sufficient to adequately capture the variability among kernels.

**Fig. 4.**
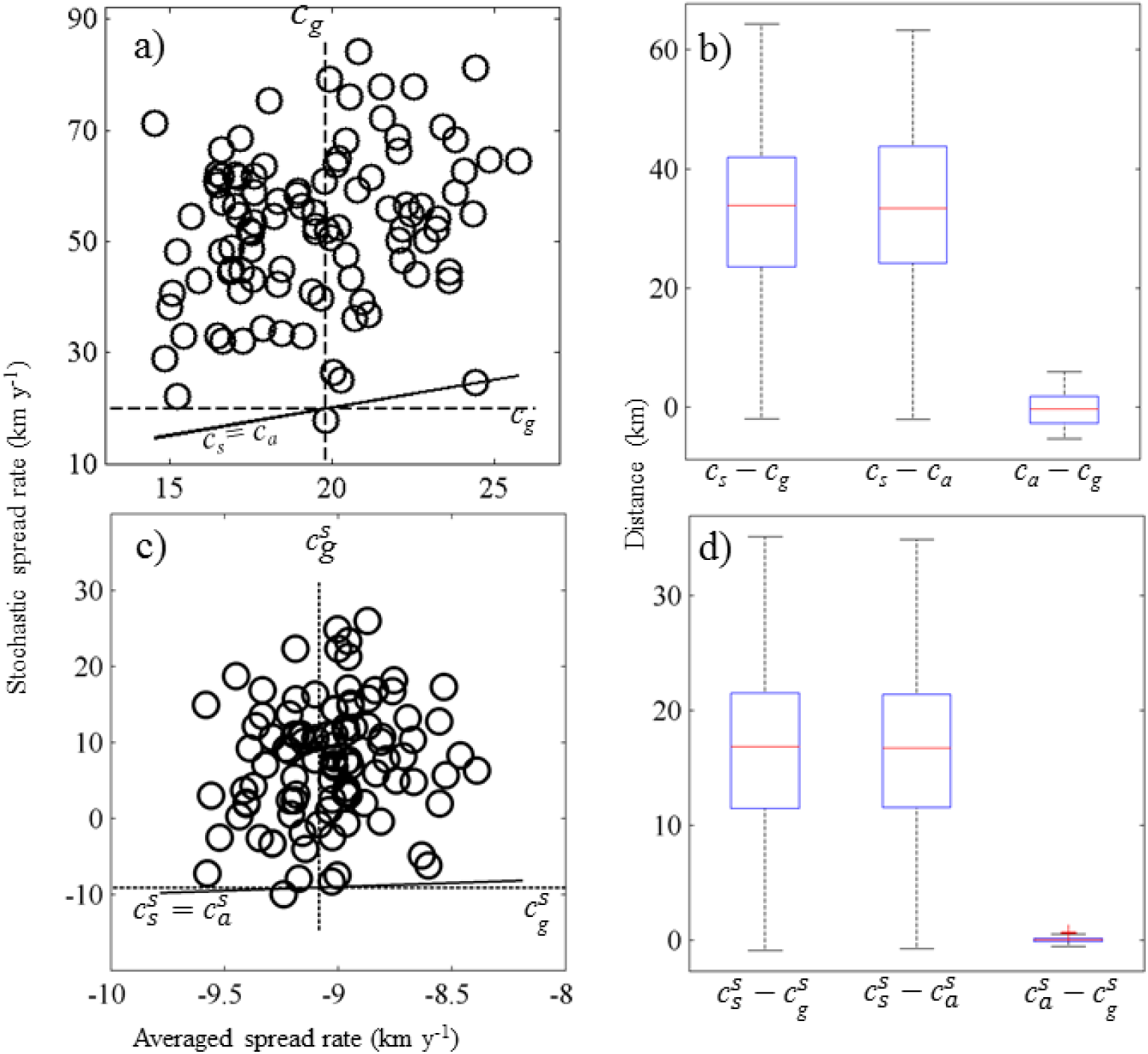
Comparison of the spread rate for stochastic, sample-averaged, and grand-averaged dispersal kernels.

Panels (a) and (b) are a scenario with only good years (featured by some northward dispersal), and panels (c) and (d) are a scenario with a mixture of good and bad years (the latter featured only southward dispersal). See text for explanation of the symbols associated with the spread rate *c*. Each point in the scatterplots in panels (a) and (c) represents a pair of simulated spread rates, averaged and stochastic, for a given random sample (a 30-year sequence of dispersal kernels or parameters; see Fig. S.2 and Table S.1). If a point is above the solid line, then the stochastic spread rate is greater than the corresponding averaged rate. The dashed line indicates the spread rate resulting from the grand-averaged dispersal kernel (Fig. S.2), which has the value *c*_*g*_ =21.5 km y^−1^ in panel (a) and 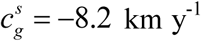 in panel (c). The box plots in panels (b) and (d) present the differences between the different types of modeled spread rates (stochastic, sample-averaged and grand-averaged). The other model parameters were fixed as *b*_A_ = 0.8, *r* = 23 adult female offspring per female per year, *T*_*1*_ = 50 d and *T*_2_ = 90 d for all simulations.

## Discussion

Analysis of our deterministic model (Eqs. 1-5) showed that spread and range expansion of marine organisms (such as aquatic invasive species) against a current can occur as a consequence of the compensatory relationship between the demographic parameter (recruitment rate, *r* = *ϕb*_*L*_*b*_1_*b*_2_), the larval diffusion coefficient (*D*) and the larval net rate of displacement (*v*). The compensatory relationship, which implies that a higher recruitment rate or a higher diffusion can compensate for the negative effect of larval advection in the direction of a current, is known to hold in many similar models and is likely a general property of spread models [3,9,18]. Spread and range expansion can also be aided by year-to-year variations in dispersal [3]. As a second step in our modeling exercise (which was the main objective of our study), time-varying larval dispersal (using a hydrodynamic model) was estimated and incorporated into the population model, thereby converting it to a stochastic model. Our results indicated that when dispersal parameters vary with time, knowledge of the time-averaged dispersal process is insufficient for determining the spread rates of the population. Specifically, and as an example, we showed that for the green crab invasion along the east coast of North America, northward spread is possible over a number of years, even when there are only a few “good years” featuring some northward dispersal among many “bad years” featuring only southward dispersal. In the following discussion, we compared our results to previous studies investigating effects of environmental stochasticity on population dynamics. We included an examination of the different ways that dispersal heterogeneity can be incorporated into a spatial model.

### Interplay between demography, dispersal and behavior

The compensatory relationship we observed between recruitment (*r*), diffusion (*D*) and displacement (*v*) appears to be robust in a broad class of spatial ecological models. In a seminal paper, Fisher derived a hyperbolic relationship between diffusion and the intrinsic growth rate of a population using a single-stage partial differential equation [18]; the relationship was extended to a model with advection by Aronson and Weinberger [37]. Pringle and coauthors [3,38] also observed a similar relationship in a stochastic cellular automaton model. In a previous paper [9], we presented an example to support the generalization of this theory to structured IDEs (i.e., with multiple stages) with simple dispersal kernels (Gaussian and Laplace). Here, we demonstrated the compensatory relationship in a structured IDE with a mechanistically motivated dispersal kernel (Eqs. 3-5).

Organisms can effectively increase *D* and decrease *v* through behavioral adaptations exploiting variations in currents. Thus, more caution should be taken when estimating dispersal parameters for organisms from oceanographic studies. For example, Bonardelli et al. [39] estimated advection in the Northumberland Strait between 2 and 8 km d^−1^ in a southeast direction; these estimates are similar to other estimates for the general Gulf of St. Lawrence region [40,41]. If this range of advection is used as an upper bound for the net rate of displacement of larvae, *v*, then our model suggests that for diffusion coefficients in the range of 10 to 100 km^2^ d^−1^, the recruitment rate, *r*, should range from 10,000 to 100,000 female recruits per female adult per year to maintain an upstream spread. This range of recruitment is not realistic for green crabs (literature suggests *r* = 23 female recruits per female adult per year; see [9]). Note that the estimate of 2-8 km d^−1^ is for passive particles drifting in surface currents. However, larvae can swim vertically to take advantage of different currents [34,42,43], which may result in a higher diffusion coefficient and/or lower net rate of displacement [44]. Indeed, to examine the effect of such behavior, we used the hydrodynamic model with vertically swimming larvae to estimate diffusion and drift parameters for the mechanistic kernel. The results led to realistic estimates of larval diffusion and, more importantly, showed high annual and spatial variability in diffusion and drift (Fig. S.2).

### Time-varying dispersal parameters in population IDEs

In addition to exploiting the vertical structure in currents, organisms may also take advantage of temporal variability in currents, such as short-term reversals in current, and turbulence (e.g., [11,12]). Studying the effect of time-varying dispersal parameters on invasion spread rates in structured models has only recently received attention of theoretical ecologists. In a stochastic cellular automaton model, Pringle and coauthors [3,38] simulated a deterministic IDE with dispersing larvae and sessile adult stages. In their simulation, larval mean advection downstream was sampled from a Normal distribution in each generation. They concluded that year-to-year variability of larval dispersal can significantly aid retention of organisms that spawn for multiple years [3]. Caswell et al. [29] provided a stochastic version of an earlier deterministic model of invasion speed for stage-structured populations [28]. For populations structured by a continuous state variable, Ellner and Schreiber [45] similarly provided a stochastic version of an earlier deterministic model for invasion speed [46]. Both Caswell et al. [29] and Ellner and Schreiber [45] found that year-to-year variability in dispersal accelerated population spread. Our work with stochastic dispersal kernels supports this, as it resulted in higher spread rates than using the corresponding time-averaged dispersal kernels.

Dispersal heterogeneity can be incorporated into population models in various ways. In our work, the dispersal heterogeneity includes both variability at the level of individuals and from one year to the next. Specifically, the hydrodynamic model that we used to simulate daily flow fields for a given year and coastal location included an individual-based particle tracking model [20,21,32,33] to obtain daily settler displacements within the larval settlement period. This is analogous to Stover et al.’s [47] “heterogeneity”, where the diffusivity of individuals is sampled from a probability distribution. They showed that intra-annual variation in dispersal leads to leptokurtic dispersal kernels, increases the population’s spread rate, and lowers the critical reproductive rate required for persistence in the face of advection. In our work, the hydrodynamic model also provided us with a set of realistic dispersal kernels (see Fig. S.2 and Table S.1 in the online supplement) that we sampled to represent year-to-year variability. This is somewhat analogous to Ellner and Schreiber’s [45] approach, which included an annual, random change in the importance (or weight) of two modes of dispersal, namely short-distance dispersal and long-distance dispersal. Other means of incorporating dispersal heterogeneity that we could explore include heterogeneity in larval behavior and development (for an example of a within year process) or randomly changing the order of “good” and “bad” years (for an example of a between year process).

Population persistence in an advective environment is theoretically related to the ability to invade against a current: aspects such as dispersal heterogeneities which affect spread rates also affect persistence in similar ways [19]. For example, Williams and Hastings [48] considered metapopulation persistence in a patchy marine coastal environment, where the unidirectional dispersal by larvae between patches is governed by a binomial random variable. This year-to-year stochasticity in ocean flow patterns increases the long-run growth rate of the metapopulation and predicts persistence, compared to using a long-term average of ocean flow. Note that an opposite result has been observed in other modeling studies on persistence in metapopulations [49,50]. For example, Watson et al. [49] showed that growth rates calculated from a constant, averaged connectivity between populations rather than the time series (connectivity versus year) were typically higher and so overestimated the likelihood of persistence. Obviously, more modeling studies are needed to better understand when heterogeneities in dispersal lead to persistence and when they do not.

### Other considerations and future work

Geometric versus arithmetic averaging of dispersal kernels has an effect on the spread rate of IDE models. Indeed, if the growth rate of a population varies over time, the total population size is described by the geometric average of annual growth rates [51]. Using the same logic, this geometric averaging also appeared in the computation of asymptotic spread rates for stochastic IDE models [29]. In our paper, we compared spread rates arising from arithmetic averages, which is a common averaging approach. Stover et al. [47] noted that the asymptotic spread rate resulting from arithmetic averaging is always higher than that from geometric averaging. Thus, had we used geometric averaging, the difference between the stochastic spread rate and that calculated from the averaged dispersal kernels would have been even larger (box plots in Fig. 4, panels b and d). Therefore, the type of averaging done in modeling or experimental studies needs to be considered carefully.

The simplification of a uniform settling rate of larvae in our model has also recently been used in modeling green crab population dynamics [52]. Other forms of probability distributions for settlement rate include Gaussian when settlement is normally distributed during the settlement period (i.e., with a main event in the middle of the period) [27], peaked when there is a one-time particularly strong settlement event at some point during the settlement period [53,54], decay when settlement is initially high and decreases over time [55], and, more generally, gamma when there is one main settlement event which could be modeled any time during the settlement period [56]. Such settlement probability distributions are useful approximations for various settlement temporal patterns and should be investigated further in the field as well as in modeling exercises. However, beyond this consideration on shape of the probability distribution for settlement rate, it is likely that dispersal and settlement should not be separated into two functions as is typically done (including in our study). The dispersal and settlement components of the kernel incorporate all sorts of physiological, behavioral and hydrodynamic processes, including larval developmental rates and behaviors (e.g., responding to differential currents, water temperatures and/or other cues) [57]. The interconnectedness among dispersal, development and settlement processes needs to be better understood and formally modeled. Our work provides a good platform from which to further refine the modeling component for larval behavior and demography (i.e., growth, development, survival, settlement) in large-scale hydrodynamic models.

Schreiber and Ryan [58] emphasized the importance of stochastic models to address predictions of increased interannual variability as a result of climate change. Our model can be used to assess the effect of environmental stochasticity on rates of spread and range expansion of marine organisms. More specifically, rising sea temperatures across the globe could lead to the appearance of anomalous flow patterns, with incidental effects on dispersal and population dynamics in many marine species [59]. Further, it has been documented that the increase in water temperature causes pelagic larval durations to be shortened and results in lower dispersal distances for marine organisms [60]. Decreased larval development times in warmer water could influence the dispersal kernel and spread rates [3]. Byers and Pringle suggested that increased temperatures would decrease spread, based solely on the shorter time spent in the water current [3]. However, the faster development may also increase settlement probability and decrease larval mortality, which may lead to increased spread rate. Our more detailed dispersal kernel (Eqs 3-5) is a first step in this direction to develop and provide tools that predict spread rates under various conditions and climate change scenarios (e.g., [61] for recent climate change scenarios). Such tools are essential to inform maps of range expansions and risk maps of areas prone to habitat degradation due to aquatic invasive species’ activities.

## Conclusion

Population spread is a complex ecological phenomenon involving the interplay between a number of processes, including demography and dispersal of organisms, and variability in these processes. Our results showed that incorporating time-varying dispersal into population models is important to enhance understanding of spatial dynamics and population spread rates. We showed that spread against a dominant current is possible even when there are only a few “good” years (featuring dispersal against the dominant current) among many “bad” years (featuring only dispersal in the dominant direction). This enhanced understanding of dispersal patterns helps guide quantifications of connectivity among populations, explain measurements of genetic structure of populations, and untangle the past history of population and community dynamics that may not correspond to observed averaged current patterns. Furthermore, improved estimation of spread rates is key to developing effective responses to biological invasions in marine systems.

## Acknowledgements

We thank the participants of the workshop on Sustainability of Aquatic Ecosystem Networks (Mathematics of Planet Earth 2013, Atlantic Association for Research in the Mathematical Sciences) for insightful discussions, as well as two anonymous reviewers for useful comments.

## Supporting Information

**Figure S.1. Release locations of particles (representing green crab larvae) in the Northumberland Strait, Canada**.

The locations are numbered from 1 to 8, from the southeast to the northwest. The dashed line is the modeled coastline used for calculating dispersal displacement of projected particles.

**Figure S.2. Simulated dispersal kernels**

Larval dispersal kernels obtained from simulations of the hydrodynamic model of the Gulf of St. Lawrence [20,21] for different locations in the Northumberland Strait (see Fig. S.1) and different years. For a given panel, the frequency distribution of the displacements of dispersing particles (representing green crab larvae) is in blue, and the fitted theoretical dispersal kernel (Eqs. 3-5) is in red. See Table S.1 for the estimated values of the pairs of *v* (net rate of displacement) and *D* (diffusion coefficient) corresponding to each simulated kernel.

**Table S.1. Estimated pairs of (*v,D*)**.

Pairs of (*v,D*) for larval dispersal kernels simulated from the hydrodynamic model for the Gulf of St. Lawrence and then estimated using the fitted Eqs. 3-5 (see also Fig. S.1 and S.2). Units of the net rate of displacement *v* and diffusion coefficient *D* are km d^−1^ and km^2^ d^−1^, respectively.

